# IRF8-driven reprogramming of the immune microenvironment enhances anti-tumor adaptive immunity and reduces immunosuppression in murine glioblastoma

**DOI:** 10.1101/2024.04.02.587608

**Authors:** Megan Montoya, Sara A. Collins, Pavlina Chuntova, Trishna S. Patel, Takahide Nejo, Akane Yamamichi, Noriyuki Kasahara, Hideho Okada

## Abstract

**Background:** Glioblastoma (GBM) has a highly immunosuppressive tumor immune microenvironment (TIME), largely mediated by myeloid-derived suppressor cells (MDSCs). Here, we utilized a retroviral replicating vector (RRV) to deliver Interferon Regulatory Factor 8 (IRF8), a master regulator of type 1 conventional dendritic cell (cDC1) development, in a syngeneic murine GBM model. We hypothesized that RRV-mediated delivery of IRF8 could “reprogram” intratumoral MDSCs into antigen-presenting cells (APCs) and thereby restore T-cell responses.

**Methods:** Effects of RRV-IRF8 on survival and tumor growth kinetics were examined in the SB28 murine GBM model. Immunophenotype was analyzed by flow cytometry and gene expression assays. We assayed functional immunosuppression and antigen presentation by *ex vivo* T-cell-myeloid co-culture.

**Results:** Mice with RRV-IRF8 pre-transduced intracerebral tumors had significantly longer survival and slower tumor growth compared to controls. RRV-IRF8 treated tumors exhibited significant enrichment of cDC1s and CD8+ T-cells. Additionally, myeloid cells derived from RRV-IRF8 tumors showed decreased expression of the immunosuppressive markers Arg1 and IDO1 and demonstrated reduced suppression of naïve T-cell proliferation in e*x vivo* co-culture, compared to controls. Furthermore, DCs from RRV-IRF8 tumors showed increased antigen presentation compared to those from control tumors. *In vivo* treatment with azidothymidine (AZT), a viral replication inhibitor, showed that IRF8 transduction in both tumor and non-tumor cells is necessary for survival benefit, associated with a reprogrammed, cDC1- and CD8 T-cell-enriched TIME.

**Conclusions:** Our results indicate that reprogramming of glioma-infiltrating myeloid cells by *in vivo* expression of IRF8 may reduce immunosuppression and enhance antigen presentation, achieving improved tumor control.

**Key points:**

1. GBM intra-tumoral myeloid cells are proliferative and targets for RRV therapy.
2. Expression of IRF8 significantly improves survival and slows tumor growth in murine GBM.
3. IRF8 expression in MDSCs reduces immunosuppression and enriches cDC1s *in vivo*.

**Importance of the study:** Recent publications have presented conflicting studies regarding the role of IRF8 in GBM. While some studies showed IRF8 as a negative prognostic factor, others demonstrated the conversion of tumor cells into DCs using IRF8. Here, we show that RRV-mediated delivery of IRF8, a clinically relevant modality, allows for transduction of both tumor and immune cells *in vivo*. We show that a significant survival effect relies heavily on the infection and modulation of both populations, and that even a modest number of reprogrammed intra-tumoral MDSCs can have a substantial impact on the immunological milieu, significantly enriching and activating cytotoxic T-cells. Further, this work reveals intra-tumoral myeloid cells as a target for other RRV-based gene therapies.

## INTRODUCTION

Glioblastoma (GBM) is the most aggressive type of primary brain tumor and has a median overall survival of ∼15 months^1,2^. The current standard-of-care treatments have not advanced significantly over the last 10 years. Although immunotherapy has led to breakthroughs in other cancers, significant success has not been demonstrated in patients suffering from primary brain tumors^3^. GBM is classified as a “cold” tumor due to being sparsely infiltrated with T-cells or active antigen-presenting cells (APCs), such as dendritic cells (DC)^4,5^. The major constituents of the intra-tumoral immune compartment are immunosuppressive myeloid cells, such as myeloid-derived suppressor cells (MDSCs)^6^. MDSCs are highly enriched in settings of chronic inflammation, and their expansion results from the recruitment of immature bone marrow-derived myeloid cells followed by their reprogramming by tumor-produced signals into an immunosuppressive population^7–9^. The accumulation of MDSCs is an adverse prognostic indicator in primary and recurrent GBM patients, underscoring the crucial need to therapeutically modulate these cells^10^. Previous studies have attempted to deplete MDSCs or inhibit their recruitment using 5-Fluorouracil, anti-CCL2 antibodies, radiotherapy, and others^11–13;^ however, these studies have not aimed to reverse the fundamental issue of immunosuppression.

Due to their undifferentiated state, MDSCs are relatively plastic and are a promising target for reprogramming^14,15^. Accordingly, as a novel approach, we hypothesized that it might be possible to convert these immature infiltrating myeloid cells into mature APC-like cells that are capable of antigen cross-presentation. To this end, we focused on interferon regulatory factor 8 (IRF8), a transcription factor and master regulator of the type 1 conventional DC (cDC1) lineage^16,17^. cDC1s exist in both lymphoid and tissue-resident states, where they excel at cross-presentation and induction of anti-tumor responses via CD8 T-cell activation. Interestingly, IRF8 has also been shown to be a negative regulator of MDSC development^18–20^. Therefore, we hypothesized that the delivery and augmentation of IRF8 in GBM-TIME could lead to the reprogramming of MDSCs toward a cDC1-like phenotype.

As a gene delivery mechanism, we employed a replicating retroviral vector (RRV), which has been used safely in clinical trials of glioma patients^21–23^. RRVs are nonlytic, result in stable genomic integration, and are only able to infect actively proliferating cells, as they do not inherently contain a nuclear localization sequence^24,25^. Furthermore, RRV infection is inhibited by innate antiviral host defenses and cleared by adaptive immunity, which act to prevent further replication in normal cells and tissues These mechanisms are impaired or suppressed in cancer cells, hence RRV replication has been shown to be highly tumor-selective. Because of these characteristics, RRV is a useful delivery system for gene therapies in the unique environment of GBM, where tumor cells, but not healthy brain tissue, are rapidly proliferating. Within tumors, each infected cancer cell becomes a new source for further viral spread and gene delivery. In this study, we evaluated the proliferative and reprogramming capability of intra-tumoral myeloid cells. We hypothesized that direct infection of myeloid cells with an RRV expressing IRF8 would mitigate immunosuppression and enhance the anti-tumor T-cell response.

## MATERIALS AND METHODS

### Replicating Retroviral Vectors

**Plasmid generation:** IRF8 sequence was taken from NCBI (Gene ID: 15900). Vectors were designed using SnapGene software suite (https://www.snapgene.com). DNA fragment assembly was done using HiFi Gibson Assembly Master Mix (Invitrogen, A46628). Resulting clones were screened using PCR and sequenced for accuracy. Confirmed clones were expanded, and plasmid DNA was isolated using a Maxiprep kit (Invitrogen, K210016). **RRV production:** RRVs were made using a standard calcium phosphate transfection as described in Supplementary Materials and Methods. Vectors were titrated using SB28 cells; flow cytometry was used to measure both Gag and P2A expression.

### SB28 glioma cell line

Details on the establishment of the SB28 cell line were previously described^26,27^ Green fluorescence protein (GFP) was knocked out in all SB28 cells used in this study. GFP expression in the parental SB28 cell line was disrupted using CRISPR, GFP-negative cells were then sorted out of the resulting pool population, expanded, and used in all further studies. Media composition is detailed in Supplementary Materials and Methods.

### SB28-OVA glioma cell line

Generation of SB28 cell line expressing full-length OVA peptide was previously described^28^. OVA sequence from Addgene (#22533) was used.

### Cell doubling time assay

SB28 WT, SB28 RRV-EMPTY (100% transduced) cells, and SB28 RRV-IRF8 (100% transduced) cells were plated at 1x10^5^ cells (n=3 per time point) and counted (Thermofisher Countess 3)at 24 and 48 hours post-seeding. Doubling times were averaged among replicates and time points.

### Secreted factor assay

SB28 WT, RRV-EMPTY 100% transduced, and RRV-IRF8 100% transduced cells were seeded at 3x10^4^ cells and cultured for 6 days. The resulting conditioned media was centrifuged, and the supernatant was filtered through a 0.7µm filter. The LEGENDplex flow cytometry-based assay (BioLegend, 740446) was used to measure secreted factor concentration according to the manufacturer’s protocol. Data were analyzed using LEGENDplex Analysis Software.

### Orthotopic glioma models

6–10-week-old female C57BL/6J mice (Jackson Laboratories) were used in all animal experiments. Animals were housed and handled in the vivarium at the University of California San Francisco. All procedures followed an approved Institutional Animal Care and Use Committee (IACUC) protocol. The procedure for orthotopic tumor inoculation is described in Supplementary Materials and Methods. Pre-mixed RRV-transduced cells were prepared as described in Supplementary Materials and Methods.

### Subcutaneous glioma model

4x10^5^ SB28 cells in 100µL of cold HBSS were mixed in 1:1 ratio with Matrigel. 200µL of cell/Matrigel slurry was injected subcutaneously in the right flank. Tumor measurements were obtained via caliper and tumor area was calculated using length (mm) x width (mm).

### Isolation of tumor-infiltrating cells

Tumor-bearing brain quadrant was isolated and manually and enzymatically dissociated. A detailed protocol description is available in Supplementary Materials and Methods

### Flow cytometry

Single-cell suspensions were labeled with fluorescent antibodies (surface and intracellular antigens) according to the protocol outlined in Supplementary Materials and Methods. A list of antibodies used is available in Supplementary Table 1.

### RNA preparation and gene expression assay

RNA was extracted from previously frozen tumor samples using RNeasy Mini Kit (Qiagen, 74104) and normalized to 100ng/µL. For gene expression assays, the Nanostring nCounter Mouse PanCancer Immune Profiling panel was used (Nanostring, XT-CSO-MIP1-12). Data were analyzed using nSolver and Rosalind software.

### 3′-Azido-3′-deoxythymidine (AZT) administration via drinking water

Mice were given AZT or 2% control sucrose water *ad lib* from 2 days pre-tumor inoculation until study endpoint. A description of AZT dose and administration is detailed in Supplementary Materials and Methods.

### Immunosuppression: Myeloid cell/T-cell co-culture

Myeloid cells from RRV-EMPTY or RRV-IRF8 tumors were isolated and co-cultured with CFSE-stained T cells from naïve non-tumor bearing C57BL/6J mice. A detailed description is in Supplementary Materials and Methods.

### Antigen presentation: DC/CD8 T-cell co-culture

DCs derived from tumors and cervical lymph nodes were isolated from OVA RRV-EMPTY or OVA RRV-IRF8 mice and co-cultured with naïve OT-1 T cells. A detailed description is in Supplementary Materials and Methods.

### Statistical Analyses

Mantel-Cox (Log-rank) test was used to determine the significance in Kaplan Meier curves (GraphPad Prism v10.1.0). For experiments comparing RRV-EMPTY versus RRV-IRF8, results were analyzed using Student’s t-test. For studies with more than two groups, results were analyzed using one-way ANOVA. Significance symbols correspond to the following: *p < 0.05; **p < 0.01; ***p < 0.001; ****p < 0.0001.

## RESULTS

### Intra-tumoral myeloid cells are proliferating and are a viable target for RRV-based **therapies**

As we aimed to target the intra-tumoral myeloid compartment, we first characterized this population in our SB28 model^26^. The SB28 model of GBM is clinically relevant, with its low mutational burden and resistance to immune checkpoint blockade therapy^27^. Like human GBM, SB28 orthotopic tumors are highly infiltrated by myeloid cells, which comprise the vast majority of intra-tumoral immune cells^28,29^ (Fig. 1a). We further evaluated the myeloid compartment based on Ly6C and Ly6G, surface markers commonly used to distinguish monocytic (M-MDSCs) and polymorphonuclear/granulocytic MDSCs (PMN-MDSCs), respectively^30^. In SB28 tumors, M-MDSCs were the dominant MDSC population (Fig. 1a), with previous studies in other models showing that M-MDSCs are more effective at suppressing T-cell functions than PMN-MDSCs^31^. Within the overall myeloid cell population, 36.6% (± 3.859 standard deviation (SD), n=6) of cells expressed Arginase 1 (Arg1), an intracellular enzyme and marker of immunosuppression^32,33^. When analyzed further, 37.1% (±4.94 SD, n=6) of CD11b+ Ly6C+ cells also highly expressed Arg1, (Fig. 1b) suggesting these M-MDSCs had an immunosuppressive phenotype. Therefore, M-MDSCs in our model represent a robust population of the TIME and are a promising target for novel myeloid-targeting therapies, such as RRV-based genetic reprogramming.

**Figure 1.**
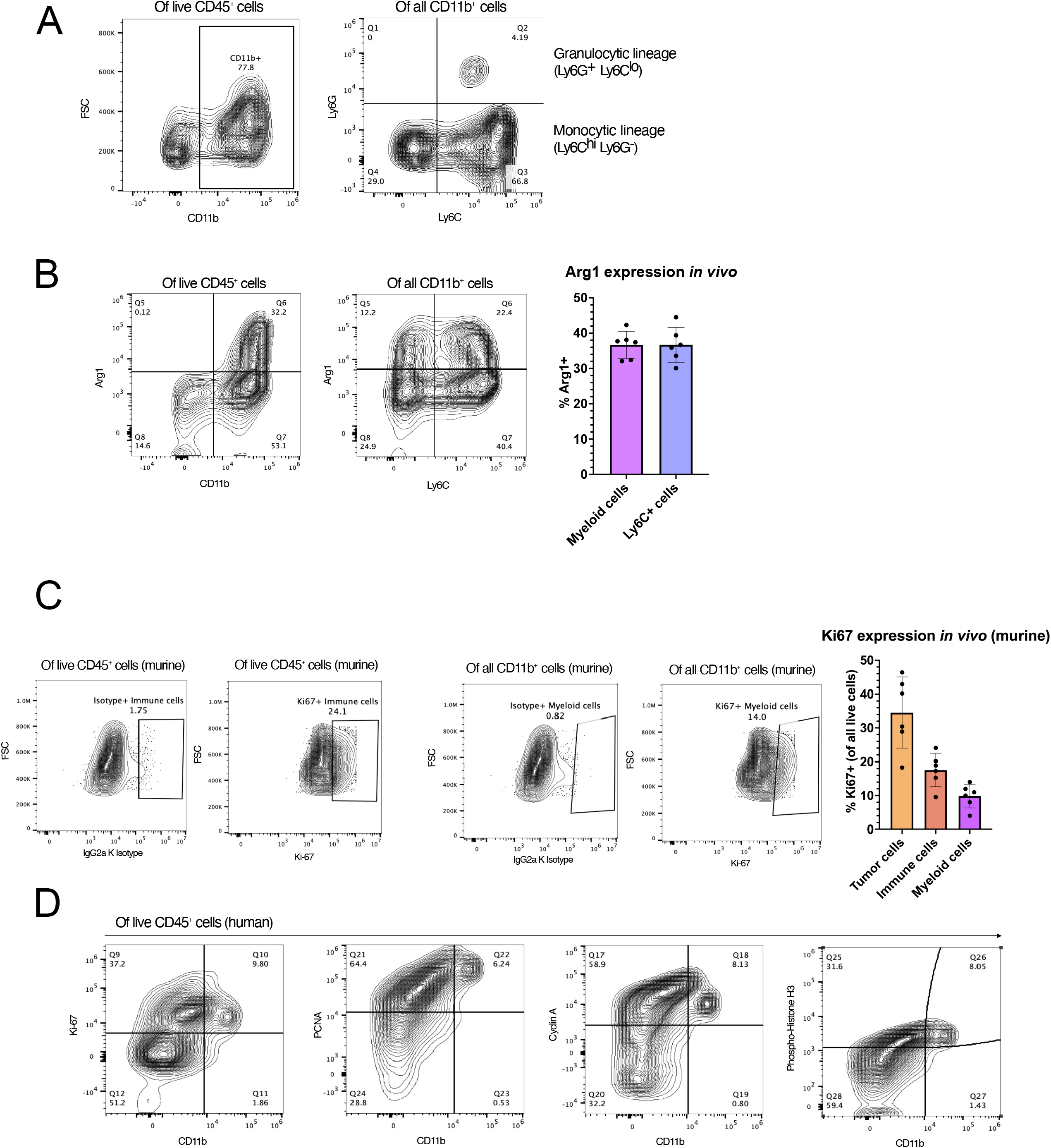
Human and murine glioblastoma contain proliferating myeloid cell populations. **(A)** Characterization of SB28 intra-tumoral myeloid cell populations. Tumors were harvested on day 18 post-tumor inoculation and analyzed by flow cytometry. The left panel shows live myeloid cells (CD45+ CD11b+); the right panel shows M-MDSCs (CD45+ CD11b+ Ly6C^hi^ Ly6G-) and PMN-MDSCs (CD45+ CD11b+ Ly6G+ Ly6C^lo^). **(B)** Flow cytometric analysis of Arg1 expression in all myeloid cells (CD45+ CD11b+) and M-MDSCs (CD45+ CD11b+ Ly6C+). Bars represent mean of 6 biological replicates. **(C)** Expression of intracellular Ki67 expression in immune (CD45+) and myeloid (CD45+ CD11b+) cells. IgG2a K isotype was used to define gates. Bar graph (right) represents *in vivo* expression of Ki67 in tumor, immune, and myeloid cells. Bars show the mean of 6 biological replicate samples. **(D)** Expression of proliferation markers Ki67, PCNA, Cyclin A, and Phosphorylated Histone H3 in human primary GBM samples. Flow cytometry plots are pre-gated for live CD45+ immune cells.

Transduction with RRV requires active cell division^24,25^. To determine whether we could target intra-tumoral myeloid cells with RRV, we examined their expression of Ki67, a proliferation marker. Within all live cells analyzed, 17.56% (± 4.92 SD, n=6) of all immune cells, and 8.83% (± 3.49 SD, n=6) of myeloid cells, were positive for Ki67 at the time of tumor harvest, day 17 post-tumor inoculation. (Fig. 1c). Because these values represent Ki67+ cells at a single time point, cumulatively a more significant number of myeloid cells will have undergone mitosis over the lifetime of the tumor. As an integrating virus, RRV is highly persistent, and intratumoral replication and cellular transduction will continue over time, so these data suggest that a portion of the myeloid compartment may be a viable target for RRV-based therapies.

To evaluate the relevance to humans, we evaluated the expanded list of proliferation markers in clinical samples obtained from patients with primary GBM (n=2 patients). We isolated tumor-infiltrating immune cells from surgically resected fresh tumor samples and evaluated the expressions of PCNA (expressed in G1 and S phases), cyclin A (late S, G2, and M phases), and phosphorylated histone H3 (p-histone H3; M phase). Interestingly, we found the majority of human CD45+CD11b+ cells were positive for all four proliferation markers (Fig. 1d).

### IRF8 transduction of SB28 tumor cells *in vitro* decreases CCL2 secretion but does not impact proliferation capacity

We inserted a transgene cassette encoding the murine transcription factor IRF8 into a Moloney murine leukemia virus (MMLV)-based RRV (RRV-IRF8)., In this vector, the P2A sequence encoding a “self-cleaving” peptide, which can be used to identify transduced cells by intracellular staining and flow cytometry, links the transgene cassette to the virus genome. The control vector encodes the P2A sequence but not IRF8 (RRV-EMPTY) (Fig 2a).

**Figure 2.**
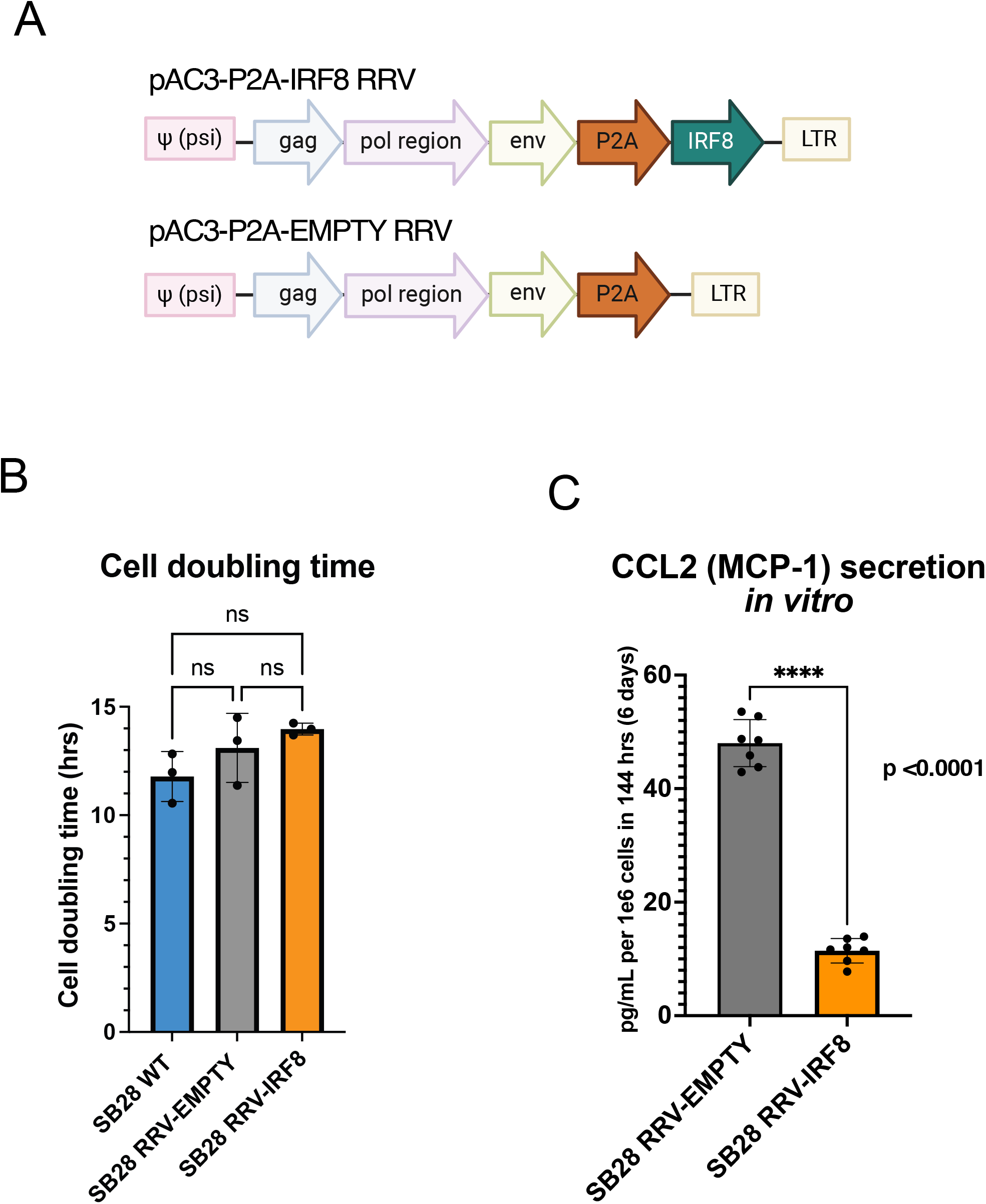
RRV-IRF8 transduction of SB28 murine glioblastoma *in vitro*. **(A)** Vector maps of RRV-IRF8 and RRV-EMPTY control. Both vectors contain the P2A self-cleaving peptide linking the transgene cassette to the viral genome. The P2A sequence is also used as a marker for vector transduction following intracellular detection and flow cytometric analysis. **(B)** Cell doubling times of SB28 WT (non-transduced), SB28 RRV-EMPTY, and SB28 RRV-IRF8. Cell were counted at 24- and 48-hours post-seeding. Doubling times were averaged among six technical replicate wells. **(C)** MCP-1/CCL-2 secretion in SB28 RRV-EMPTY versus SB28 RRV-IRF8 cells *in vitro*. Cells were cultured for 6 days before conditioned media collection; media was filtered to exclude debris and cell components.

We first evaluated whether infecting SB28 cells *in vitro* with the RRV-IRF8 or RRV-EMPTY vectors might have any effect on cell growth. Compared to untransduced SB28 WT-cells, there was no significant effect on cell doubling time after transduction with either the RRV-EMPTY or RRV-IRF8 vectors (Fig. 2b), implying that neither vector transduction per se, nor exogenous expression of the IRF8 transgene impacted tumor cell proliferation rate. To better understand the effect of IRF8 transduction in GBM cells, we characterized the *in vitro* tumor cell culture conditioned media, using a flow cytometry-based secretome assay. Of note, the assay revealed that, among the tested 13 chemokines, the secretion of CCL-2 (MCP-1) was most prominently downregulated as a result of IRF8 transduction (Fig. 2c). CCL-2 is a major chemoattractant in GBM that recruits effector and regulatory T-cells (Tregs) and immature myeloid cells to the tumor^34^. In patient samples, elevated CCL2 is correlated with worse patient outcomes, and inhibition of CCL2 in mouse models reduces intra-tumoral MDSCs and increases T-cell cytotoxicity^12,35,36^. Interestingly, intra-tumoral myeloid cells from RRV-IRF8 transduced SB28 tumor-bearing mice *in vivo* showed significantly decreased expression of the CCR2 receptor (Fig. S1a).

### Transduction with IRF8 suppresses the growth of intracerebral SB28 GBM tumors

To evaluate the effects of IRF8 expression *in vivo*, we tested the RRV-EMPTY and RRV-IRF8 using a pre-mixed tumor establishment model, in which 2% of the tumor cells implanted were transduced with either the RRV-IRF8 vector or RRV-EMPTY control vector. This allows for a single intracranial injection procedure, reducing inflammation and disruption of the blood-brain barrier associated with multiple injections and survival surgeries. Pre-mixing the RRV at a low percentage allows for efficient and reproducible tumor inoculation and enables the RRV to efficiently initiate replication and spread immediately following tumor engraftment. Tumor growth kinetics were monitored with bioluminescent imaging (BLI), and tissues were harvested at a scheduled timepoint or humane endpoint (Fig 3a).

**Figure 3.**
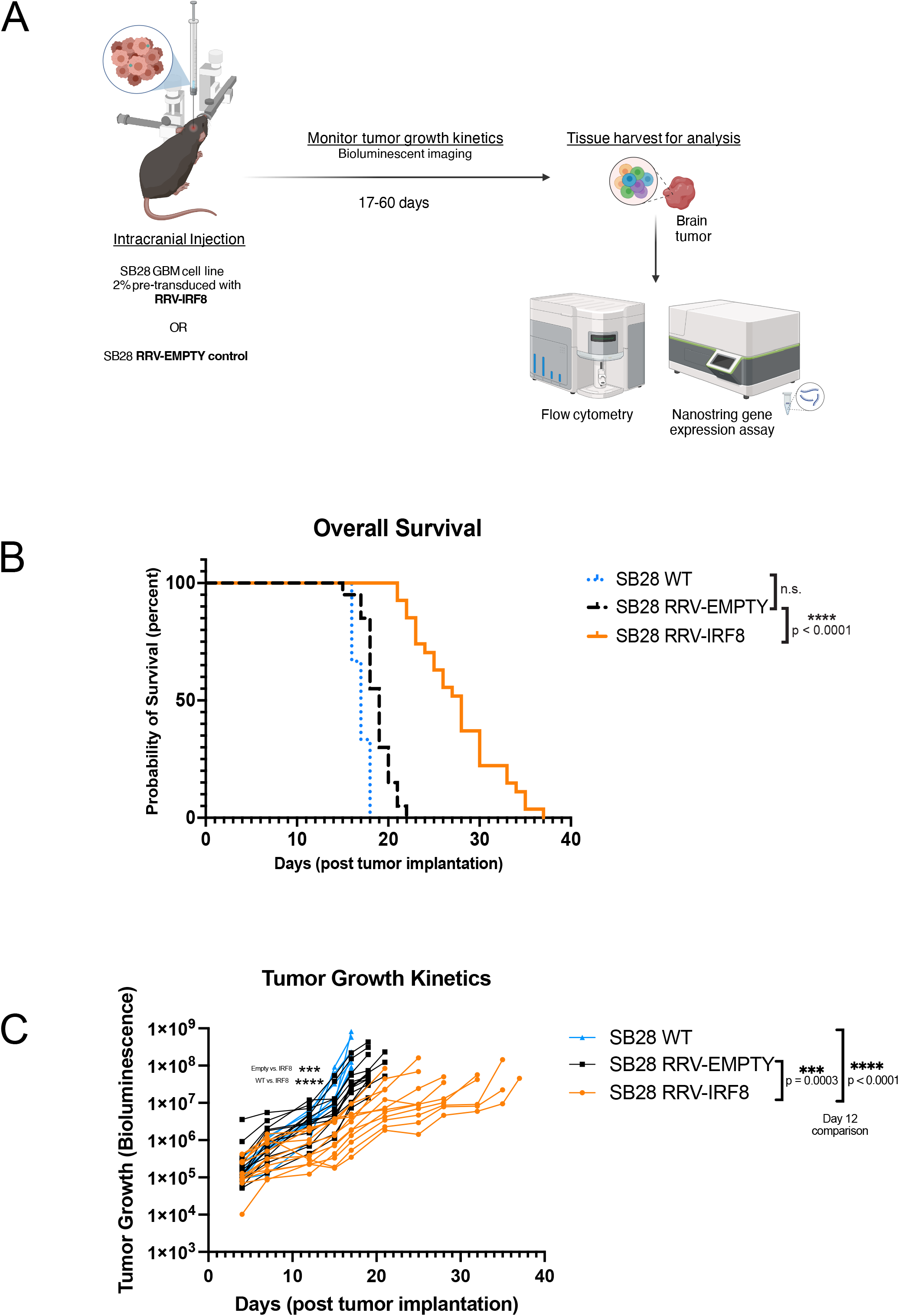
Transduction with IRF8 *in vivo* suppresses the growth of intracerebral SB28 tumors. **(A)** Schematic of *in vivo* studies. SB28 cells pre-transduced at 2% with either RRV-EMPTY or RRV-IRF8 were implanted intracerebrally. Tumor growth kinetics were monitored using bioluminescence (BLI) twice per week until study completion. Tissues were harvested and dissociated into single cells for analysis. **(B)** Kaplan-Meier curves showing survival; SB28 WT, SB28 RRV-EMPTY, and SB28 RRV-IRF8, **(C)** BLI imaging data corresponding with tumor growth kinetics. P-values assessed on day 12 post-tumor inoculation.

Mice with SB28 RRV-IRF8 tumors had significantly longer overall survival (median overall survival (mOS)=28 days) than either untransduced SB28 WT (mOS=17 days, p<0.0001) or RRV-EMPTY (mOS=19 days, p<0.0001) groups (Fig. 3b). BLI revealed that RRV-IRF8 mice also had slower growth kinetics than SB28 WT and RRV-EMPTY groups (p<0.0001, day 12 post-tumor inoculation). RRV-EMPTY and RRV-IRF8 tumors grew at similar rates until approximately day 12, when the two groups began to separate (median luminescence 5.3x10^5^ vs. 3.6x10^6^, p = 0.0003) and remained lower for the duration of the study (Fig. 3c).

### IRF8 transduction enhances the number of GBM-infiltrating T-cells and type 1 conventional dendritic cells

Because of the significant survival benefit and tumor growth delay observed *in vivo*, in the absence of cell proliferation rate change *in vitro*, we hypothesized that overexpression of intra-tumoral IRF8 led to an overall change in the TIME, perhaps associated with reduced CCL2. Nanostring analysis of overall gene expression in bulk SB28 tumors from each treatment group showed clear segregation of RRV-EMPTY and RRV-IRF8 transduced tumors into two distinct groups (Fig. 4a). Tumors transduced with RRV-IRF8 showed a higher abundance of CD45+ cells when compared with controls. The abundance of different immune cell types, each defined by a subset of genes (Fig 4b, right box), was given a score, and the scores for the control (RRV-EMPTY) and experimental (RRV-IRF8) groups were plotted on the left in Fig. 4b. Consistent with Fig. 4a, IRF8 expression resulted in the overall increase in many immune populations, especially T-cells and cytotoxic cells.

**Figure 4.**
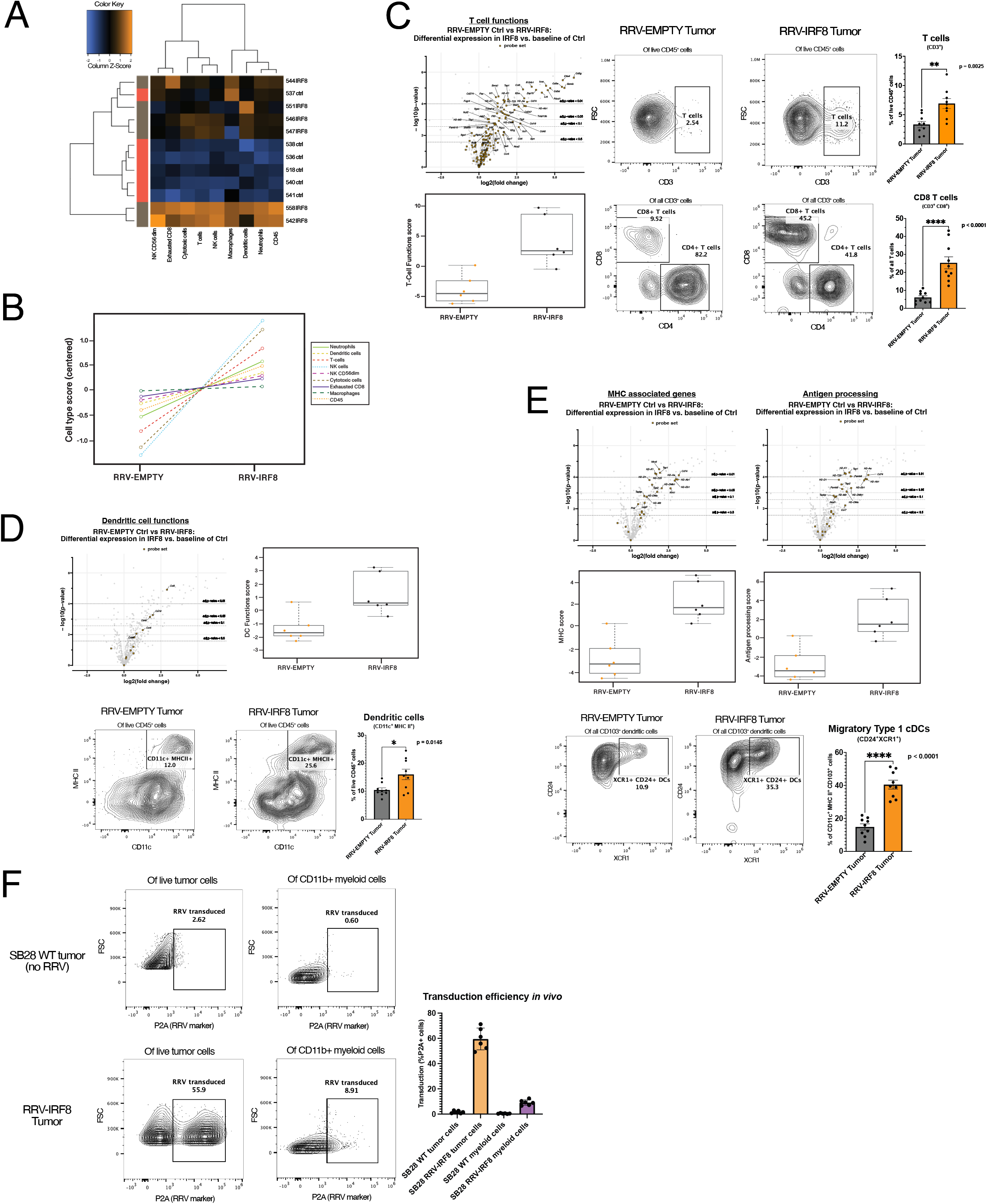
IRF8 transduction enhances the number of glioblastoma-infiltrating T-cells and type 1 conventional dendritic cells. **(A)** Heatmap of differential expression of immune cell types between RRV-EMPTY (red bars, n=6 biological replicates), and RRV-IRF8 groups (grey bars, n=6 biological replicates). Total RNA was isolated from day 18 tumors. **(B)** Immune cell type changes between groups. Each cell type is associated with a set of genes; differential expression of gene sets is correlated with cell type abundance. Cell type profiling algorithm was previously described by Danaher *et al* (PMID: 28239471). **(C)** Left panel, volcano and box-and-whisker plots derived from the expression of T-cell genes (Supp. Table 2). In volcano plots, circular dots represent all differentially expressed genes; T-cell genes are represented with squares. Dashed horizontal lines correspond with adjusted p-value cut-offs. Right, representative flow plots of pan T-cells (CD45+ CD3+) and CD4 (CD45+ CD3+ CD4+) or CD8 (CD45+ CD3+ CD8+) T-cells. Bars represent the mean of 9 biological replicates. **(D)** Volcano and box-and-whisker plots derived from the expression of DC genes (Supp. Table 3). Representative flow plots of the pan-DC population (CD45+ CD11c+ MHC II+). Bars represent the mean of 9 biological replicates. **(E)** Volcano and box-and-whisker plots derived from the expression of MHC-associated or antigen-processing genes (Supp. Table 4). Bottom panels show flow cytometric analyses of CD103+ DCs derived from day 18 tumors. Live cells were gated on CD45+ CD11b-CD11c+ MHC II+ and CD103. The cDC1 populations were further refined by selecting CD24+ XCR1+, markers for terminally differentiated cDC1s capable of antigen cross-presentation. Bars represent the mean of 9 biological replicates. **(F)** Representative flow plots of *in vivo* RRV transduction using P2A as the marker for transduced cells. Top panels are SB28 WT (no RRV; non-transduced) tumors; the bottom panels are RRV-IRF8 tumors. Bars represent mean of 6 biological replicates.

Next, we evaluated the impacts of IRF8 expression on the T-cell compartment of the TIME, using a T-cell targeted Nanostring gene expression assay and flow cytometry. As illustrated in the volcano plot (Fig 4c, top left), Nanostring analysis showed that in RRV-IRF8 tumors, 38 T-cell-associated genes were significantly differentially expressed (see also Supp. Table 2). Interestingly, IRF8 transduction most significantly upregulated the levels of *CD3g, Ctla4,* and *Gzmb,* suggesting that IRF8 expression enhanced the infiltration of activated and cytotoxic T-cells, with an overall enhancement of T-cell functionality (Fig. 4c, bottom left). Flow cytometric analyses detected a significant (p=0.0025) increase of T-cells in RRV-IRF8 tumors compared to control (RRV-EMPTY) tumors (Fig. 4c, top right). On evaluation of the CD4 and CD8 T-cell compartments, we observed that CD4 cells were the majority of T-cells in control tumors, comprising approximately 80% of total T-cells. However, in IRF8-RRV tumors, the CD8 T-cell population was significantly enriched (p < 0.0001) with CD4 T-cells comprising only about 42% of total T-cells (Fig. 4c, bottom right).

We hypothesized that while innate and adaptive immune mechanisms restricted RRV infection and spread in normal tissues, the permissive tumor microenvironment would allow for infection of proliferating intratumoral myeloid cells. Thus, any immature myeloid cells transduced by RRV-IRF8 could adapt a more cDC1-like phenotype. Indeed, as illustrated in the volcano plot in Fig. 4d, Nanostring analysis of the intra-tumoral DC compartment shows upregulation of 6 genes associated with DC functions in RRV-IRF8 tumors as compared to controls (see also Supp. Table 3). In support of this data, flow cytometry analyses revealed an enrichment of the pan-DC population (CD11c+ MHC II+) in RRV-IRF8-transduced tumors (p = 0.015, Fig. 4d). Further gene expression analysis revealed significant upregulation of genes associated with MHC class I (*H2-Aa, H2-Ab1*), MHC class II (*H2-Eb1*), and antigen processing and presentation (*Tap1, Psmb9*), among others (Fig. 4e, top). Further flow cytometric analyses showed significant enrichment of the cDC1 (CD11c+, MHC II+, CD103+, CD24+, XCR1+) population in RRV-IRF8 tumors, compared to controls (p < 0.0001; Fig 4e, bottom), suggesting that enhanced infiltration by cytotoxic T-cell and cross-presenting cDC1s contribute to the survival benefit and delayed tumor growth kinetics observed in Fig. 3.

To further elucidate the effects of exogenous IRF8 expression in the TIME, we measured RRV transduction in tumor cells versus myeloid cells. We observed that approximately 59.52% (± 8.62 SD, n=6), of tumor cells were RRV-positive. More intriguingly, although lower than tumor cells, about 9.11% (± 2.14 SD, n=6), of the myeloid cell populations were also RRV-positive, indicating successful spreading and transduction of RRV into proliferating myeloid cells. A review of recent literature indicated that a small number of mature DCs can provide critical anti-tumoral functions in the immunological milieu and even a modest increase in APCs abundance can improve the anti-tumor immune response^37^. As RRV was observed to spread to over half of the tumor cells starting from the initial 2% pre-transduced SB28 cell inoculum, we did not disregard the contribution of IRF8 expression from either population and sought to investigate this intriguing phenotype further.

### Infection of intra-tumoral immune cells by RRV-IRF8 is necessary for decreased tumor growth rate and survival benefit

We aimed to answer a vital mechanistic question: whether the transduction of tumor cells alone is sufficient to cause the observed TIME changes and survival benefit, or if the modest population of transduced myeloid cells contributes in an essential manner. To this end, we designed a study where we compared experimental conditions under which (1) only tumor cells were infected and RRV spread is restricted or (2) all proliferating cells could be infected and the RRV is allowed to spread freely as in previous studies. To achieve this, the first group was given the anti-retroviral drug azidothymidine (AZT), a thymidine analog that inhibits reverse transcriptase and therefore precludes the ability of the virus to replicate^38^. AZT is used clinically and has been previously shown to inhibit RRV spread in mice when administration through drinking water^39^.

As a proof of concept, we evaluated the efficacy of AZT water using tumors pre-mixed with 2% green fluorescent protein (GFP)-RRV. In this model, RRV spread was quantified based on the GFP signal at day 17 from tumor inoculation. In the control group, approximately 83.90% (± 5.30 SD, n=3) of the tumor cells were GFP-positive. In contrast, in the AZT-treated group, GFP positivity was suppressed to only 1.55% (± 0.82 SD, n=3) confirming the *in vivo* efficacy of AZT (Fig. 5a). We then used the same AZT-administration scheme in our RRV-IRF8 model (Fig. S3a-experimental scheme). We stratified tumor-bearing mice into three groups (n=10) for each RRV (EMPTY and IRF8). In group 1, mice were injected with SB28 cells containing a 2% pre-mixed RRV-population with no AZT administration, recapitulating the conditions from previous studies in Fig. 3 and 4. Groups 2 and 3 were implanted with 30% and 100% pre-mixed RRV, respectively and received AZT. In these groups, RRV reverse transcription was blocked, preventing any further spread to proliferating cells, thereby restricting transgene expression solely to RRV-IRF8 already integrated into the genomes of pre-transduced tumor cells and their progeny *only.* AZT administration alone did not impact tumor growth (RRV-EMPTY/control water mOS= 17 days vs. RRV-EMPTY/AZT mOS=17.5 days). As shown in Fig. 5b-c, pre-mixed SB28 tumors established with RRV-IRF8 30% and 100% provide a modest survival benefit, with median survival times of 20.5 and 23 days, respectively (Fig. 5e, Fig. S3b). Strikingly, mice with 2% RRV-IRF8 pre-mixed tumors in which RRV spread was permitted to spread freely throughout the tumor, including immune cells, showed a significant survival benefit (mOS = 33.5 days) (Fig 5d-e) compared to the other groups, including the 100% RRV-IRF8 group + AZT. This study indicates that direct infection of non-tumor cells with RRV-IRF8 is crucial for the survival benefit and suggests that even a modest level of myeloid cell transduction (Fig. 4f) may be adequate to achieve this result.

**Figure 5.**
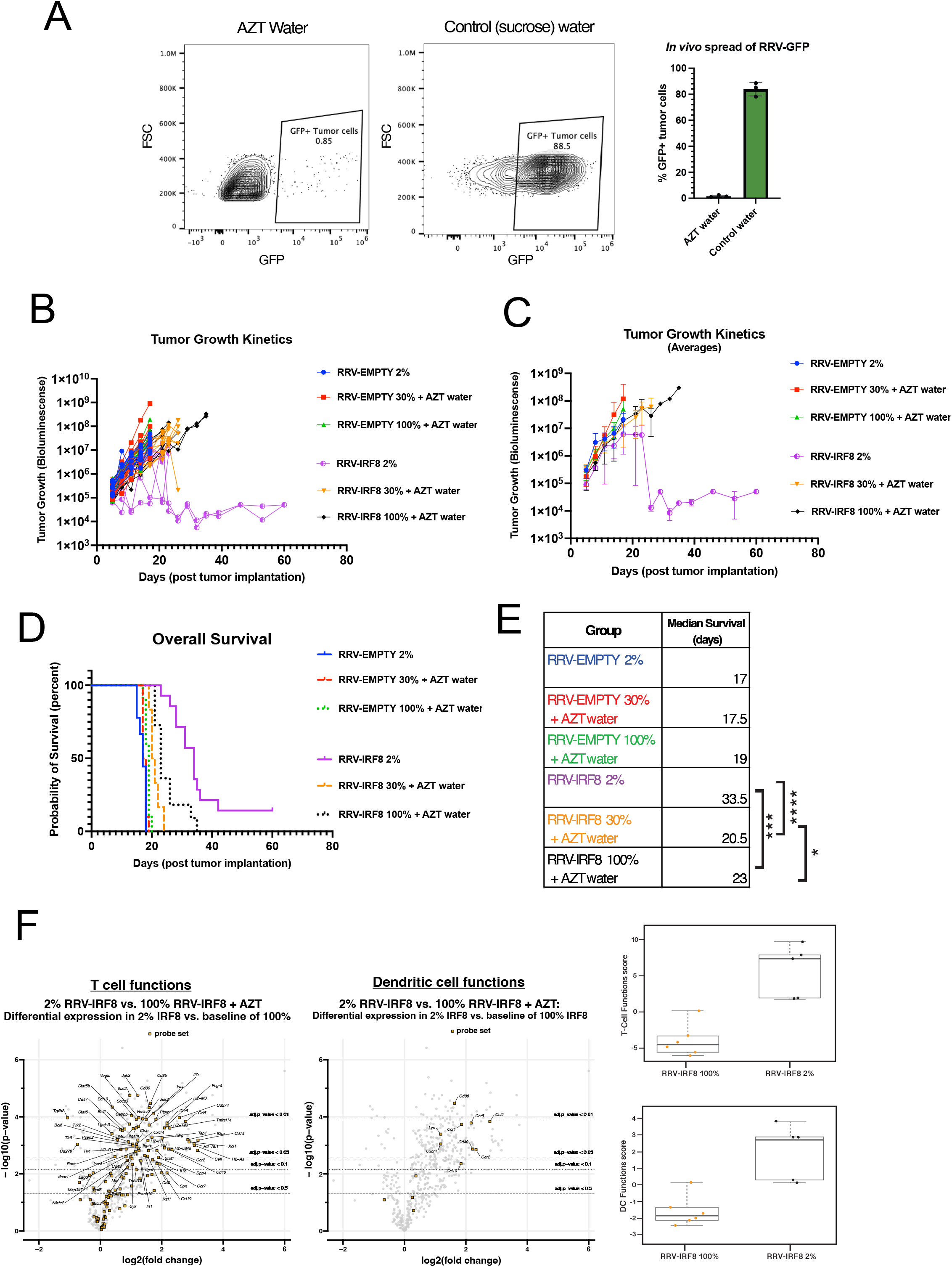
Infection of non-tumor cells by RRV-IRF8 is necessary for survival benefit and slowed tumor growth. **(A)** Mice were given 0.4mg/mL AZT + 2% sucrose water or 2% sucrose water-only control, with drug administration beginning two days prior to tumor inoculation and continuing until study endpoint (day 17 post-tumor inoculation). Representative flow plots of GFP+ tumor cells in mice receiving AZT or control water. Bars represent the mean of 3 biological replicates. **(B)** BLI tumor growth kinetics plots. 6 groups; n=10 mice per group. BLI performed twice weekly until study endpoint. BLI concluded at day 60 for 2 long-term surviving animals. **(C)** Average BLI tumor growth kinetics. **(D)** Kaplan-Meier curves showing survival. **(E)** Median survival for all groups and significance comparisons for RRV-IRF8 2% vs. 100%, RRV-IRF8 2% vs 30%, and RRV-IRF8 30% vs 100%. **(F)** Volcano plots and box-and-whisker plots show differential expression of T-cell function- and DC function-related genes between RRV-IRF8 100% + AZT (n=6 biological replicates) and RRV-IRF8 2% groups (n=5 biological replicates).

To further understand the impacts of IRF8 transduction in non-tumor cells, we used Nanostring to compare bulk gene expression in RRV-IRF8 2% pre-mixed tumors versus RRV-IRF8 100% pre-mixed tumors + AZT. RNA was isolated from tumor-bearing brain quadrant at day 17 post-inoculation. We observed differential expression of many T-cell-related genes (Fig. 5f), including some that were not seen in previous analyses (Fig. 4). Interestingly, expression of *Tgfb2* and checkpoint molecules *Cd276* (*B7-h3*) and *Lag3*, which have been identified as negative prognostic factors in GBM patients^40–42^ were downregulated in samples from the RRV-IRF8 2% group. Importantly, GBM tumors produce high levels of TGFβ2, and TGFβ signaling contributes to immunosuppression and tumor progression^43^. Furthermore, the DC compartment showed a significant upregulation of *Cd86*, a marker of mature DCs capable of activating T-cells through interaction with CD28. These data further suggest that active intratumoral replication of RRV-IRF8, associated with IRF8 transduction in myeloid cells, significantly improves anti-tumor immune responses and even reduces the expression of known GBM-promoting genes.

Two mice in the RRV-IRF8 2% group survived over 60 days post-tumor inoculation without disease progression. To assess whether these mice developed immunological memory, we rechallenged them with a subcutaneous injection of untransduced SB28 WT cells in the right flank using naïve, age-matched mice as controls (Fig. S3c). While the subcutaneous tumors grew in control mice, the rechallenged mice rejected the tumor. In summary, these data collectively suggest that additional IRF8 transduction in myeloid cells suppresses tumor-intrinsic immunosuppressive factors and enhances anti-tumor immunity, leading to the acquisition of long-term adaptive immune responses.

### RRV-IRF8 functionally reduces myeloid-derived immunosuppression and enhances antigen presentation

We sought to further characterize the functions of IRF8-reprogrammed myeloid cells. Using flow cytometry, we found intra-tumoral myeloid cells (M-MDSCs, PMN-MDSCs, and Macrophages) in SB28 RRV-IRF8 2% tumors expressed lower levels of two immunosuppressive markers, Arg1 and IDO, compared to tumors transduced with RRV-Empty (Fig. 6a-d).

**Figure 6.**
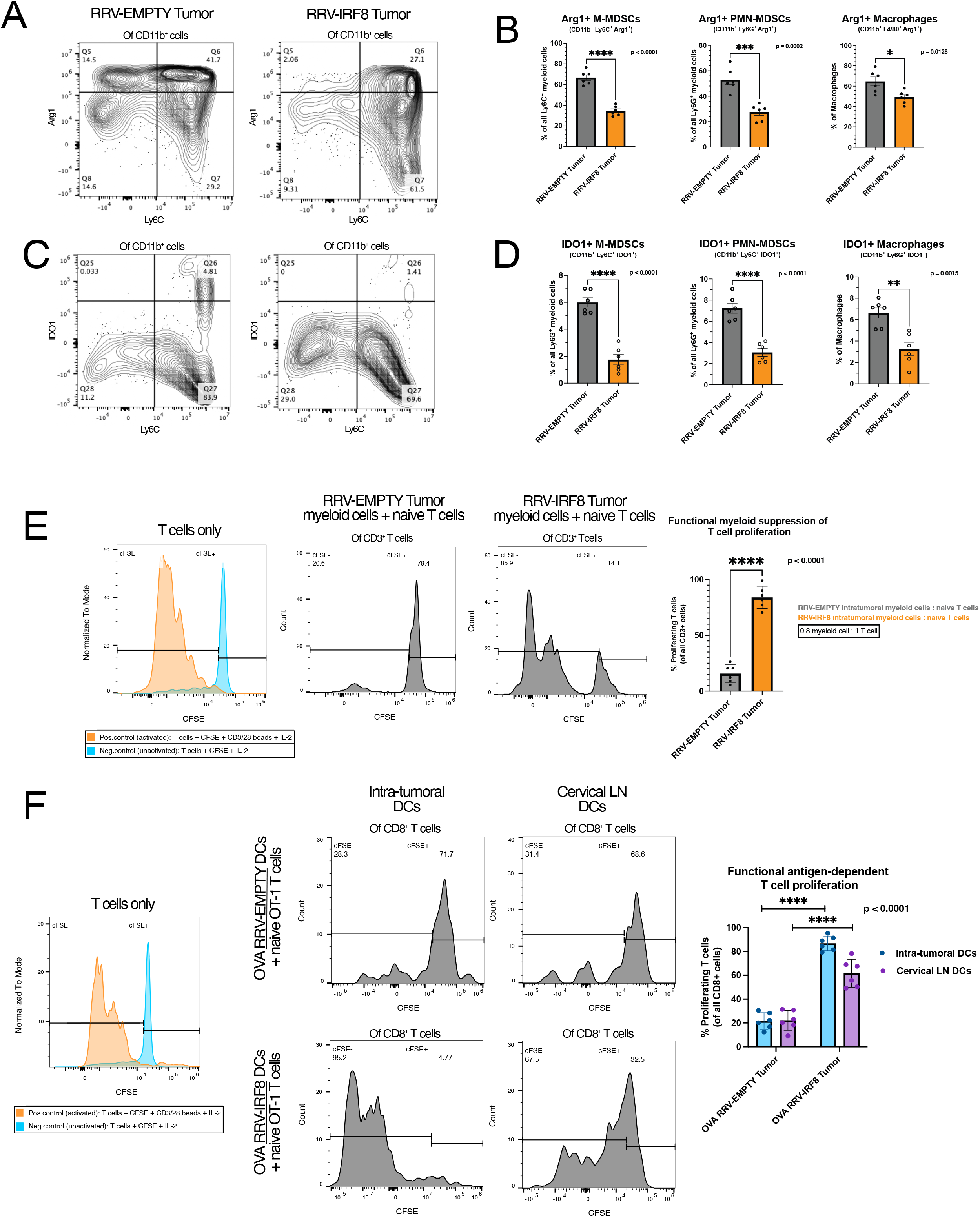
IRF8 transduction reduces immunosuppressive myeloid cells and enhances antigen presentation. **(A)** Representative flow plots of Arg1 expression in Ly6C+ cells, all plots pre-gated on live CD45+ CD11b+ cells. **(B)** Bars show Arg1 expression in M-MDSCs, PMN-MDSCs, and Macrophages, representing the mean of 6 biological replicates. **(C)** Representative flow plots of IDO expression in Ly6C+ cells, all plots pre-gated on live CCD45+ CD11b+ cells. **(D)** Bars show IDO expression in M-MDSCs, PMN-MDSCs, and F4/80+ macrophages, representing the mean of 6 biological replicates. **(E)** Left, positive and negative controls for T-cell activation; gates set on negative control peak. T-cell/myeloid cell co-culture at 0.8 effector: 1 target ratio. Intra-tumoral myeloid cells were isolated from day 18 RRV-EMPTY or RRV-IRF8 tumors. Naïve T-cells were isolated from age-matched non-tumor bearing mice. Representative flow plots show T-cell proliferation (CFSE peaks) after 4 days of co-culture. Bars represent the mean of 6 biological replicates (n=3 technical replicates for each). **(F)** Left, positive and negative controls for T-cell activation. OT-1 T-cell/DC co-culture: CD11c+ DCs were isolated from SB28 OVA RRV-EMPTY or RRV-IRF8 tumors and cervical lymph nodes. Representative flow plots show T-cell proliferation (CFSE peaks) after 4 days of co-culture. Bars represent the mean of 6 biological replicates (n=2 technical replicates for each biological replicate).

Next, to investigate the immunosuppression capabilities of myeloid cells from SB28 2% pre-mixed RRV-EMPTY versus 2% pre-mixed RRV-IRF8 tumors, we utilized a myeloid/T-cell co-culture assay. Animals from both groups were euthanized at day 17-post tumor inoculation and intra-tumoral myeloid cells were isolated. Concurrently, T-cells from age-matched naïve animals were isolated. Subsequently, myeloid cells were co-cultured with CFSE-labeled T-cells with CD3/CD28-stimulation for 4.5 days. T-cells cultured with myeloid cells from RRV-IRF8 tumors proliferated significantly more, undergoing 3-4 proliferation cycles, while T-cells cultured with myeloid cells derived from control tumors underwent 0-1 proliferation cycles (p < 0.0001) (Fig. 6e). These data suggest that IRF8 expression can functionally reprogram intra-tumoral myeloid cells to reduce their immunosuppression and to facilitate T-cell proliferation.

Finally, we tested the ability of DCs from RRV-IRF8 treated mice to activate T-cells in an antigen-specific manner. We used the ovalbumin (OVA) model antigen system, and inoculated mice with intracerebral SB28-OVA 2% pre-mixed RRV-EMPTY or 2% pre-mixed RRV-IRF8 tumors. We euthanized all animals on day 26 and isolated CD11c+ DCs from tumors and cervical lymph nodes (cLN). DCs were co-cultured with CFSE-labeled naïve OT-1 CD8 T-cells. Both intra-tumoral and cLN DCs from RRV-IRF8 treated mice induced high levels of OT-1 T-cell proliferation compared to DCs from RRV-EMPTY mice (p < 0.0001) (Fig. 6f). Interestingly, intratumoral DCs from RRV-IRF8 transduced tumors induced the most robust T-cell proliferation (∼5 proliferation cycles), suggesting that RRV-driven reprogramming induces the development of functional APCs *in situ*, after which APCs migrate to cLNs and prime T-cells.

## DISCUSSION

Lack of functional APCs and the negative contribution of immunosuppressive myeloid cells are well-recognized barriers for developing effective immunotherapy approaches for patients. A productive anti-tumor immune response relies on the crosstalk between APCs and T-cells within the tumor microenvironment^44^. We evaluated *in situ* transduction of myeloid cells with IRF8, a critical transcriptional regulator of cDC1s and a suppressor of MDSCs^16–18^ Our data indicate that RRV-mediated reprogramming of intra-tumoral myeloid cells into cDC1-like cells can lead to reduced immunosuppression and enhanced antigen presentation in the immunologically cold GBM TIME, associated with prolonged survival.

The relevance of murine GBM models remains an essential and challenging consideration when designing immune-based therapeutics. Although the SB28 orthotopic model mimics the immunosuppressive TIME of human GBM^28^, there are known differences in the MDSC compartment. Our data concur with previous reports demonstrating that the M-MDSC population is dominant in mouse tumors, while the PMN-MDSC population is dominant in human GBM^10,45^. Nevertheless, our data demonstrates that reprogramming using RRV-IRF8 directly impacts both MDSC populations, in which both Arg1 and IDO expression were significantly reduced. Interestingly, Trovato *et al.*^46^ and Groth *et al.*^31^ reported that M-MDSCs are more immunosuppressive than PMN-MDSCs and have higher capacity to inhibit T-cell proliferation. This suggests that, although M-MDSCs may be present in lower absolute numbers in GBM patients, their contribution to immunosuppression can be significant, and, therefore, reprogramming this subset cells may represent a promising therapeutic modality. Our approach, which efficiently reverts immunosuppression in both MDSC subsets and promotes antigen presentation, would likely improve anti-tumor immunity in patients as well.

While our data implicated a critical contribution of IRF8 transduction in non-tumor cells, the effects of IRF8 expression in tumor cells remains to be fully elucidated. In 2021, Gangoso *et al*. demonstrated that GBM stem cells evaded immune attack by adopting a myeloid-like transcriptional signature, including expression of IRF8^47^ and a clinical study from Lei *et al.* reported IRF8 as a negative prognostic biomarker in bulk glioma tissues^48^. On the other hand, a 2023 study by Zimmermannova *et al.* revealed an alternative role of IRF8, demonstrating that exogenous expression of IRF8 and other DC-regulatory genes directly converted tumor cell lines into cDC1-like cells, capable of processing and presenting antigens^49^. As both tumor and myeloid cells were transduced with IRF8 in our RRV system, the significance of exogenous IRF8 expression in SB28 cells must be considered. SB28 cells normally show undetectable levels of IRF8 (Fig. S1b), and RRV-IRF8 transduced SB28 cells did not demonstrate cDC1-phenotype based on expression of the DC markers MHC II, XCR1, and CD103 (Fig. S1c). These observations are consistent with the lack of gene expression changes linked to anti-tumor effects (Fig. 5f) and the only modest improvement of overall survival (Fig. 5d-e) of mice when RRV-mediated IRF8 transduction was limited to only tumor cells. On the other hand, endogenous IRF8 expression in intra-tumoral myeloid cells was varied, with IRF8 levels being inversely correlated with Arg1 expression (Fig S1b). These data suggest that IRF8 overexpression beyond its endogenous levels is required for reprogramming MDSCs into functional cDC1s.

As noted, our results demonstrate a significant impact of RRV-mediated IRF8 transduction on the immune landscape and the survival of mice bearing intracerebral SB28 tumors, even despite the modest transduction efficiency in non-tumor cells. These data further prompt us to consider the potential impacts of paracrine effects by tumor cells transduced with RRV-IRF8. *In vitro*, we showed a significant reduction of CCL2 secretion by transduced tumor cells (Fig. 2c). The CCL2-CCR2 axis recruits immature myeloid cells to the tumor, where they subsequently develop into MDSCs^50^. Notably, CCR2+ M-MDSCs have been shown to inhibit CD8 T-cell infiltration to the TIME^51^. RRV-IRF8 transduced tumors showed reduced percentages of CCR2+ myeloid cells (p=0.0464, n=6) *in vivo* (Fig. S1a). Thus, reduced CCL2 may also contribute to the observed effects, representing a paracrine role of the current RRV-mediated IRF8 transduction approach.

Another consideration is the direct transduction of myeloid cells, and whether this transduction is critical for cDC1 enrichment. While we functionally demonstrated that transduction of non-tumor cells is linked to for survival benefit (Fig. 5), it is crucial to determine whether MDSCs are truly being infected *in situ*. To this end, we examined the intracellular levels of P2A (a component of our RRV vector) by flow cytometry in both intra-tumoral cDC1s and their pan-DC counterparts and observed significantly higher levels of P2A among cDC1s, suggesting that these cDC1s were once MDSCs that were transduced and then adopted a cDC1 phenotype (Fig. S2b). The importance of antigen cross-presentation by cDC1s in potentiating anti-tumor immunity is well-reviewed in the literature, and multiple studies have shown that even a modest increase in intra-tumoral cDC1s can significantly enhance T-cell mediating tumor killing^37,52,53^

Although increased generation of cDC1s through RRV transduction is promising, ultimately, anti-tumor immunity relies also on the contribution of effector cells, including T-cells. CD4 T-cells represent the majority of T-cells in RRV-EMPTY tumors, albeit in modest absolute numbers. Examination of RRV-IRF8 tumors showed not only an increase in T-cell abundance overall, but also a shift from CD4 to CD8 T-cell dominance. Within the CD4 T-cell compartment exist Tregs, which have known immunosuppressive functions. Interestingly, some T-cells (including Tregs) can be recruited to the brain by CCL2, independent of CCR2, revealing another important role of CCL2 reduction in our approach^34,54^. Furthermore, M-MDSCs can promote Treg generation by secreting TGFß2, which was significantly downregulated in RRV-IRF8 transduced tumors^55^ (Fig. 5f). We saw a modest decrease of Tregs in RRV-IRF8 tumors (p=0.0498, n=6) compared to controls suggesting that the reduction of CD4 T-cells is due, in part, to reduced CCL2 leading to less recruitment of the Treg population (Fig. S2a). cDC1s efficiently cross-present intra-tumoral tumor antigens to CD8 T-cells, as we demonstrated in Fig. 6f, using the OVA/OT-1 model system. To further elucidate the importance of CD8 T-cells in this context, subsequent studies may utilize *in vivo* CD8 T-cell depletion. Altogether, our study demonstrates a multi-faceted impact on the recruitment of both CD4 and CD8 T-cells, concurrently reducing Treg-mediated suppression and enhancing CD8 T-cell activation.

Many challenges remain in designing and implementing immunotherapies for GBM; however, our novel gene therapy-based reprogramming approach may be a valuable tool as a primary viral-based modality or in combination with other therapies. For example, our approach presents the opportunity to combine RRV-IRF8 with CAR-T-based therapies to support the activation and persistence of engineered T-cells *in vivo*. Additionally, these studies open a new area of RRV-based gene therapies in which tumor cells are not the sole target, and RRVs may be further engineered to target myeloid cells or other populations using cell and receptor-specific promoters.

## Supplementary Materials and Methods

### Calcium phosphate transfection

293T cells were plated on Poly-L-Lysine coated dishes one day prior to transfection ddH_2_O, plasmid DNA and 2.5 M CaCl_2_ were mixed and added dropwise to 2X HBS (pH 7.12), while gently vortexing. The resulting DNA/CaPO_4_ solution was added dropwise to cells and swirled gently. The following morning, media was replaced and supplemented with 20mM HEPES and 10mM Sodium Butyrate. 5-6 hours later, the media was replaced and supplemented with 10mM HEPES. The following day, the viral supernatant media was collected and filtered through 0.45 µM syringe filter, aliquoted, and frozen at -80C.

### Media preparation

Complete RPMI (cRPMI) media was used for all cell culture: RPMI media with 10% FBS, 1% Sodium Pyruvate (final conc. 1mM), 1X MEM NEAA, 1X Glutamax, 1% HEPES (final conc. 0.01M), 1% Pen-Strep, and 0.1% Betamercaptoethanol.

### Flow cytometry

Single-cell suspensions (0.5-1x10^6^ cells/sample) of dissociated SB28 tumors were incubated with anti-CD16/CD32 Fc block (BioLegend, 156603) for 10 min, followed by viability staining (BioLegend, 423101) in PBS for 15 min. After washing, a cocktail of fluorophore-conjugated antibodies and monocyte blocker (BioLegend, 426101) was added to cells in a total volume of 100µL staining buffer (1X PBS, 0.5% BSA, 2mM EDTA) and incubated in the dark at 4°C for 30 min, rocking. For intracellular staining (cytosolic and nuclear), cells were subsequently fixed and permeabilized following the manufacturer’s protocol (FoxP3/Transcription Factor Staining Buffer Set, Invitrogen, 00-5523-00). Fluorophore-conjugated intracellular antibodies were added and incubated in the dark for at least 30 min, rocking. Samples were washed twice and suspended in 100µL staining buffer. All flow cytometry experiments were performed on the Invitrogen Attune NxT (Thermo Fisher Scientific) flow cytometer and analyzed using FlowJo software (FLOWJO, LLC, Ashland, OR, USA). All antibodies used are listed in Supp. Table 1.

### Orthotopic glioma model

Under anesthesia, mice received stereotactic tumor inoculation with 1x10^4^ cells in 2 µL HBSS (for SB28 OVA model: 2x10^4^ cells in 2 µL HBSS) at the following coordinates relative to bregma: mediolateral 2mm, dorsoventral -3mm. Mice were monitored daily and given post-operative care, per the approved IACUC protocol. Tumor growth was measured using bioluminescent imaging twice weekly: 3mg (100µL) D-Luciferin was injected intraperitoneally 10 minutes prior to image acquisition.

### Preparation of SB28-premixed cells for intracerebral injection

For each premixed cell solution, two sets of cells were prepared, SB28 WT and SB28-RRV (EMPTY or IRF8). For RRV-transduced cells, previously frozen RRV stocks were added to low-passage SB28 WT cells and allowed to spread until 100% of cells were transduced. Transduction was measured using flow cytometry staining for P2A and/or IRF8. For intracerebral implantation, SB28 WT and SB28-RRV (EMPTY or IRF8) were counted and mixed at 98% SB28 WT and 2% SB28-RRV in cold HBSS.

### Isolation of tumor-infiltrating cells

Tumor-bearing brain quadrant was dissected and manually dissociated into ∼1mm^3^ pieces. Tumor pieces were resuspended in 0.6-1 mL collagenase buffer (3.2 mg/ml Collagenase IV, 1 mg/ml Deoxyribonuclease I in PBS) and left shaking at 700 RPM at 37°C for 45 min, pausing to mix thoroughly every 15 min. Resulting dissociated tumor suspensions were filtered through a 70 µm cell strainer and washed with excess PBS; red blood cells were lysed (Lonza, BP10-548E), and cell suspensions were stored at - 80C in Bambanker (Bulldog Bio, BB01) or stained immediately for flow cytometry. Both human and mouse GBM tumors were prepared as above.

### 3′-Azido-3′-deoxythymidine (AZT) administration via drinking water

0.4 mg/mL AZT (Sigma, A2169) and 2% sucrose (Thermofisher, J65148.36) were dissolved in water and provided *ad lib* in a water bottle protected from light. As vehicle control, 2% sucrose only was used; fresh solutions were prepared weekly. To monitor water consumption, water bottles were weighed daily. Mice in the AZT/sucrose groups consumed water at the same rate as those in the control, sucrose-only groups.

### Immunosuppression: Myeloid cell/T-cell co-culture

**T-cells:** T-cells were isolated from spleens of naïve non-tumor bearing C57BL/6J mice using a CD3 bead isolation kit (BioLegend, 480023). T-cells were resuspended in 0.5 mM CFSE dye (CellTrace CFSE Cell Proliferation Kit, Thermofisher, C34570) in PBS and incubated in the dark for 10 minutes. Cells were washed several times to remove any unbound CFSE dye and were resuspended in growth medium containing with CD3/CD28 activating beads (Gibco, 11161D) and supplemented with 50 IU/mL hIL-2. **Myeloid cells:** SB28 tumors were dissociated into single cells, as described above. Myeloid cells were isolated using a CD11b bead isolation kit (BioLegend, 480109) and resuspended in cRPMI. **Co-culture:** Myeloid cells and T-cells were combined at an effector: target ratio of 0.8:1. Cells were co-cultured in cRPMI for 4.5 days and stained for flow cytometry.

### Antigen presentation: DC/CD8 T-cell co-culture

**T-cells:** T-cells were isolated from spleens of OT-1 transgenic (Jackson Laboratory, strain 003831) naïve non-tumor bearing mice using a CD8 bead isolation kit (BioLegend, 480007) and stained with CFSE dye (as above). Positive control T-cells were activated with CD3/CD28 beads, all T-cells were supplemented with 50 IU/mL **DCs:** DCs were isolated from both tumors and lymph nodes. SB28 OVA RRV-EMPTY or RRV-IRF8 tumors were dissociated into single cells as described above. Cervical lymph nodes (cLNs) from the same tumor-bearing mice were incubated with collagenase buffer for 15 min at 37C, then mechanically dissociated through a 70 µm filter to generate a single cell suspension. DCs were isolated using a CD11c bead isolation kit (Milentyi, 130-100-875). **Co-culture:** 5x10^3^ DCs were combined with 1x10^5^ OT-1 T-cells in cRPMI a 96-well plate, incubated for 4 days, and stained for flow cytometry.

## Acknowledgments

We thank the following for their contributions: Mary Helen Barcellos-Hoff and Joseph F. Costello for advice through the thesis committee; Joanna J. Phillips, Anny Shai, and the UCSF Brain Tumor Center for providing human glioblastoma tissues; Mylinh Bernardi of the Gladstone Genomics Core for assistance with RNA preparation; UCSF Core Facilities: Flow Core, Preclinical Therapeutics Core, Genome Analysis Core; members of the HO and NK laboratories for technical assistance, troubleshooting, and advice. Figures 2a, 3a, and S3a were made using Biorender.com

## AUTHORSHIP

Conceptualization: MM, HO, NK, SAC

Methodology: MM, SAC, PC, TSP, TN, AY

Investigation: MM, HO, NK

Funding acquisition: HO, NK

Project administration: HO, NK

Supervision: HO, NK

Writing – original draft: MM, HO

Writing – subsequent drafts, review, and editing: MM, HO, NK, SAC, TN, PC, TSP, AY

---

**Supplementary Figure 1.**
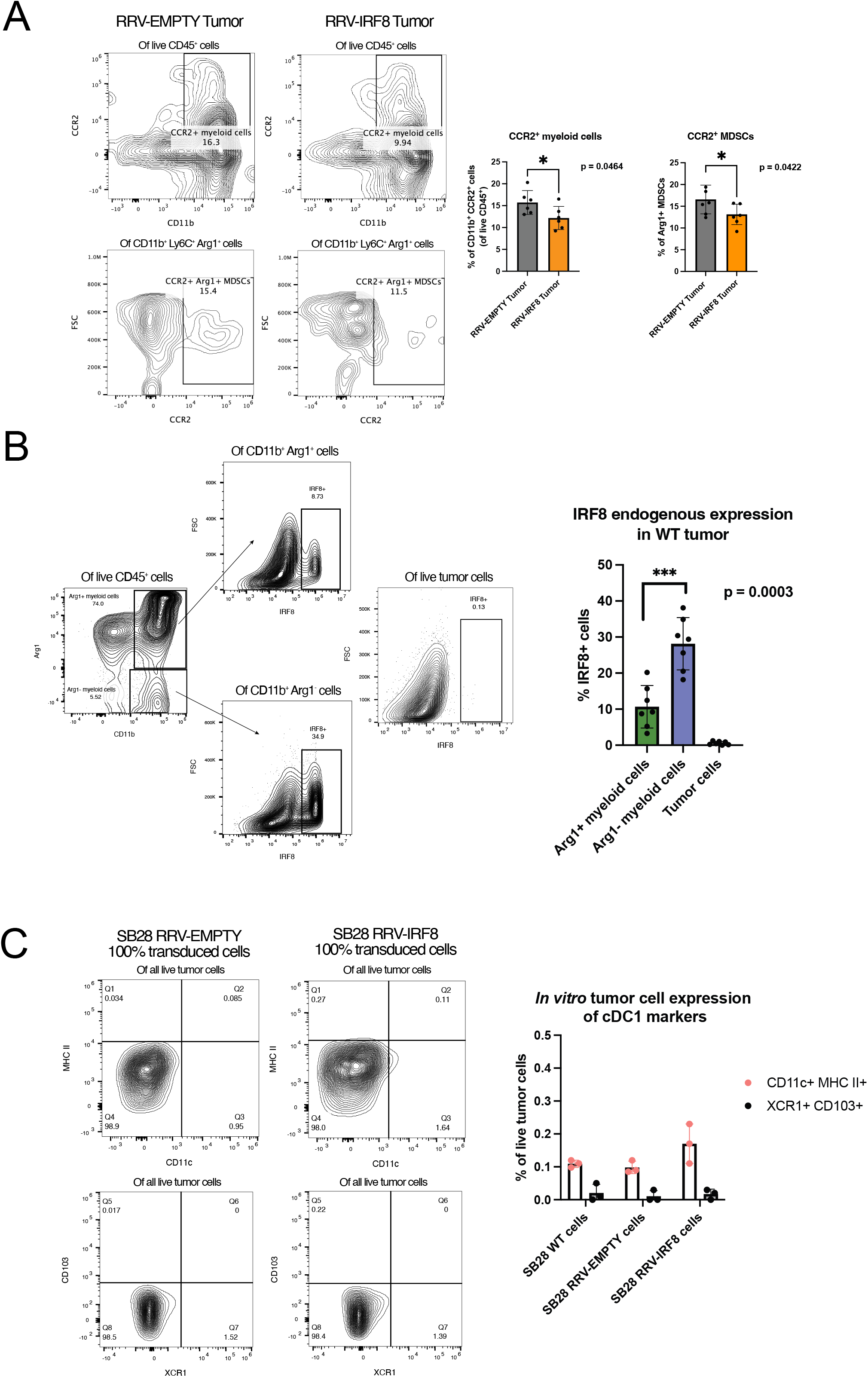
**(A)** CCR2 expression in myeloid (CD11b+) cells and M-MDSCs (Ly6C+ Arg1+) from intracerebral SB28 RRV-EMPTY or RRV-IRF8 tumors. Bars represent the mean of 6 biological replicates **(B)** Endogenous IRF8 expression in CD11b+ Arg1+, CD11b+ Arg1-, and tumor cells in intracerebral SB28 WT tumors. Bars represent the mean of 6 biological replicates. **(C)** *In vitro* expression of cDC1-associated markers in SB28 WT, SB28 RRV-EMPTY 100% transduced, and SB28 RRV-IRF8 100% transduced cell lines. Bars represent the mean of 3 technical replicates.

**Supplementary Figure 2.**
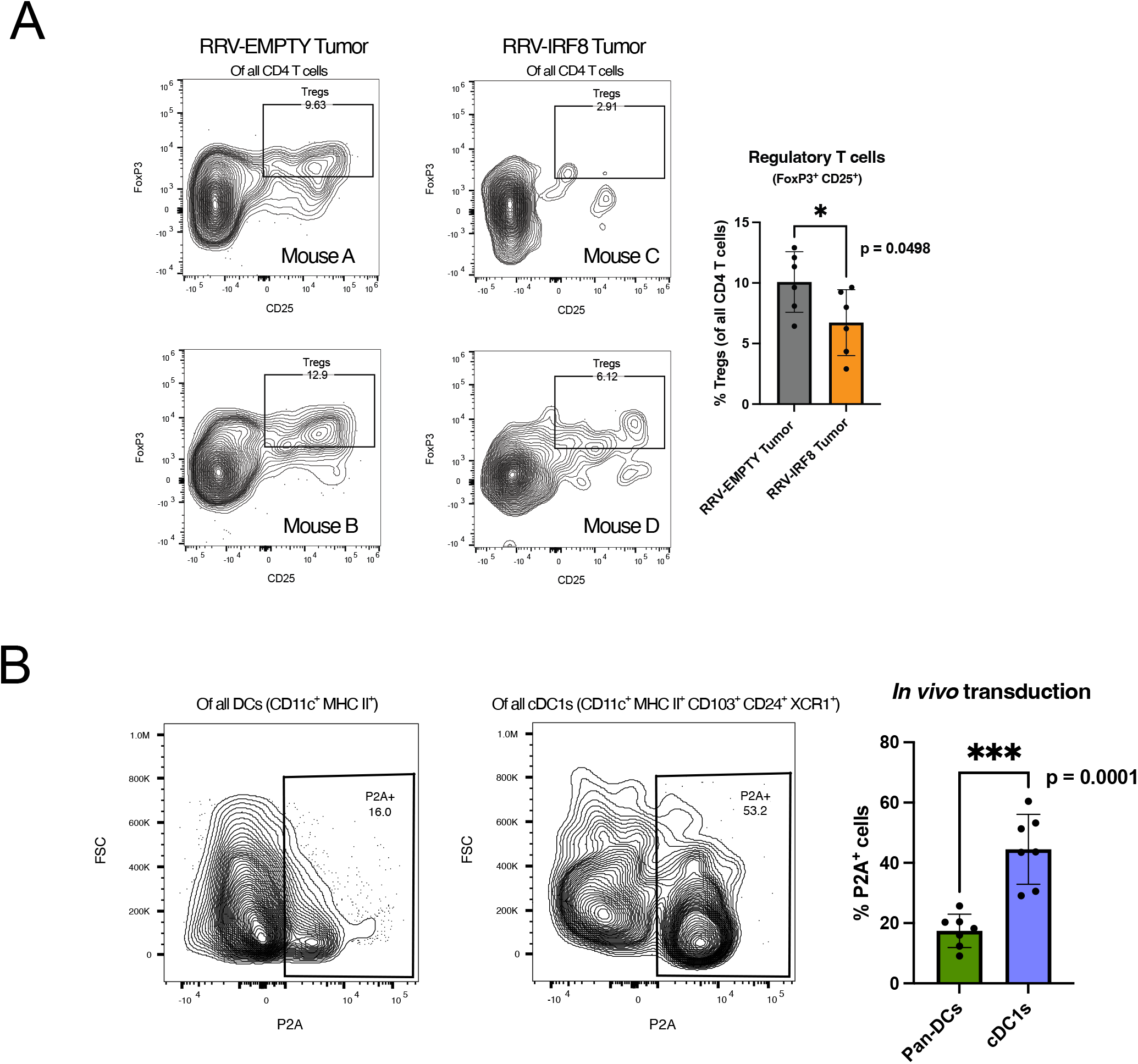
**(A)** T regulatory cell infiltration in SB28 RRV-EMPTY or RRV-IRF8 tumors. Bars represent the mean of 6 biological replicates. **(B)** *In vivo* transduction efficiency of pan-DCs (CD11c+ MHC II+) or cDC1s (CD11c+ MHC II+ CD103+ CD24+ XCR1+). Bars represent the mean of 7 biological replicates.

**Supplementary Figure 3.**
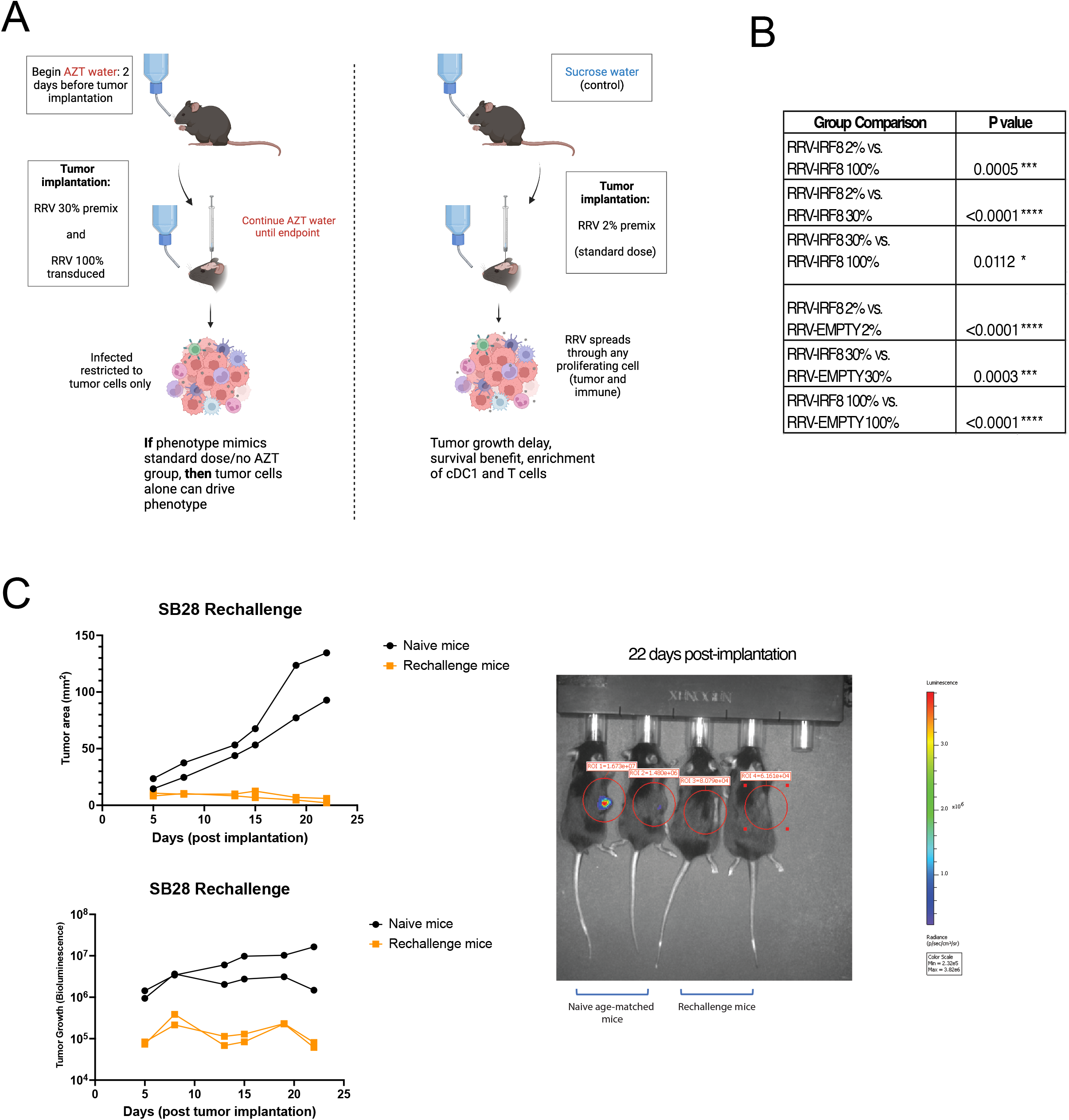
**(A)** Experimental schema for the data set presented in Fig. 5. **(B)** Group comparison statistics for Fig. 5e. **(C)** Long-term survivors from Fig. 5b-c were rechallenged with 4x10^5^ SB28 WT cells in the right flank on day 65 post-intracerebral tumor inoculation. Graphs represent tumor area (mm^2^) and tumor growth bioluminescence until day 22 post-tumor inoculation.

**Supplementary Table 1:**
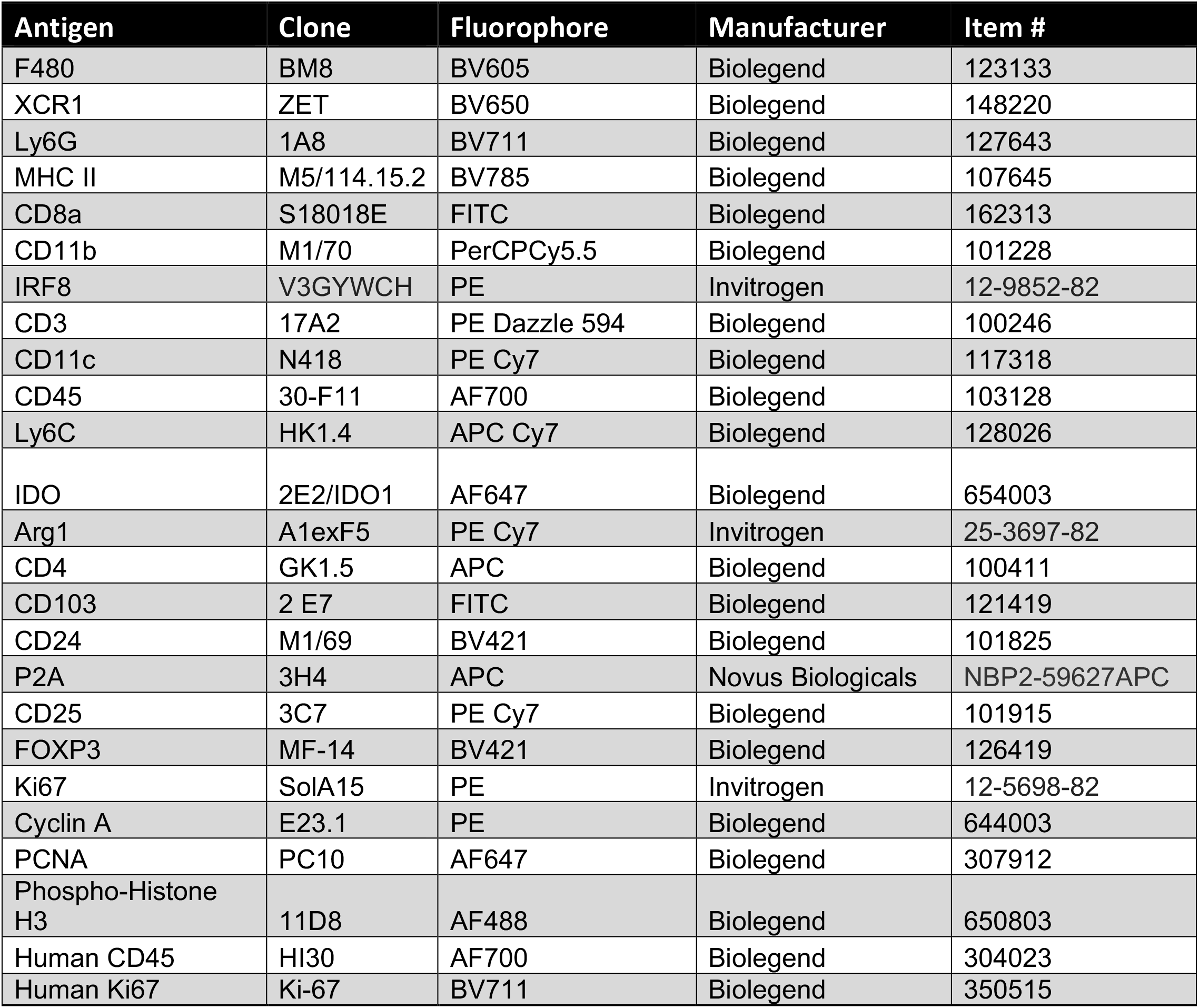
Antibodies used in flow cytometry.

**Supplementary Table 2:**
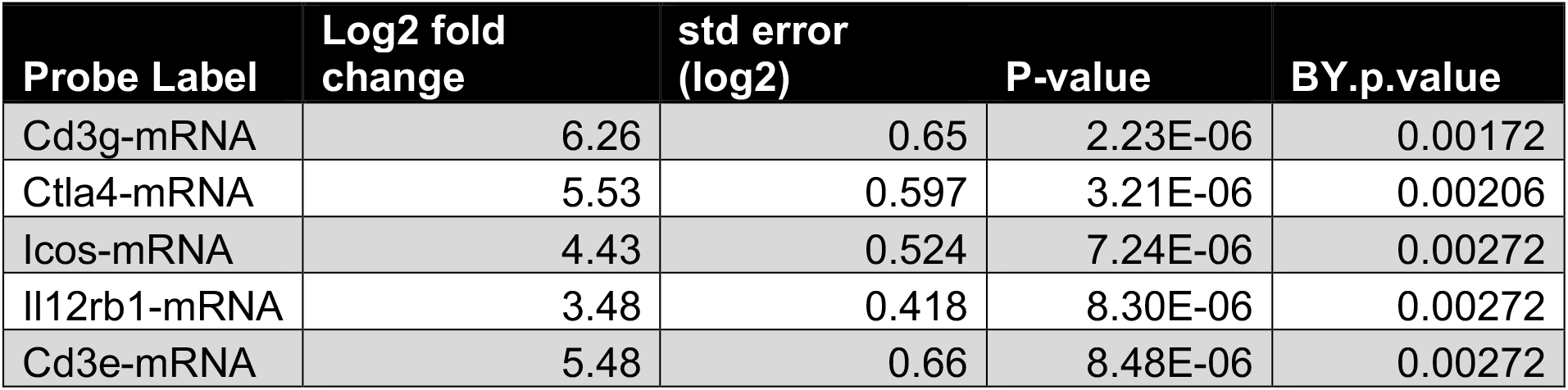
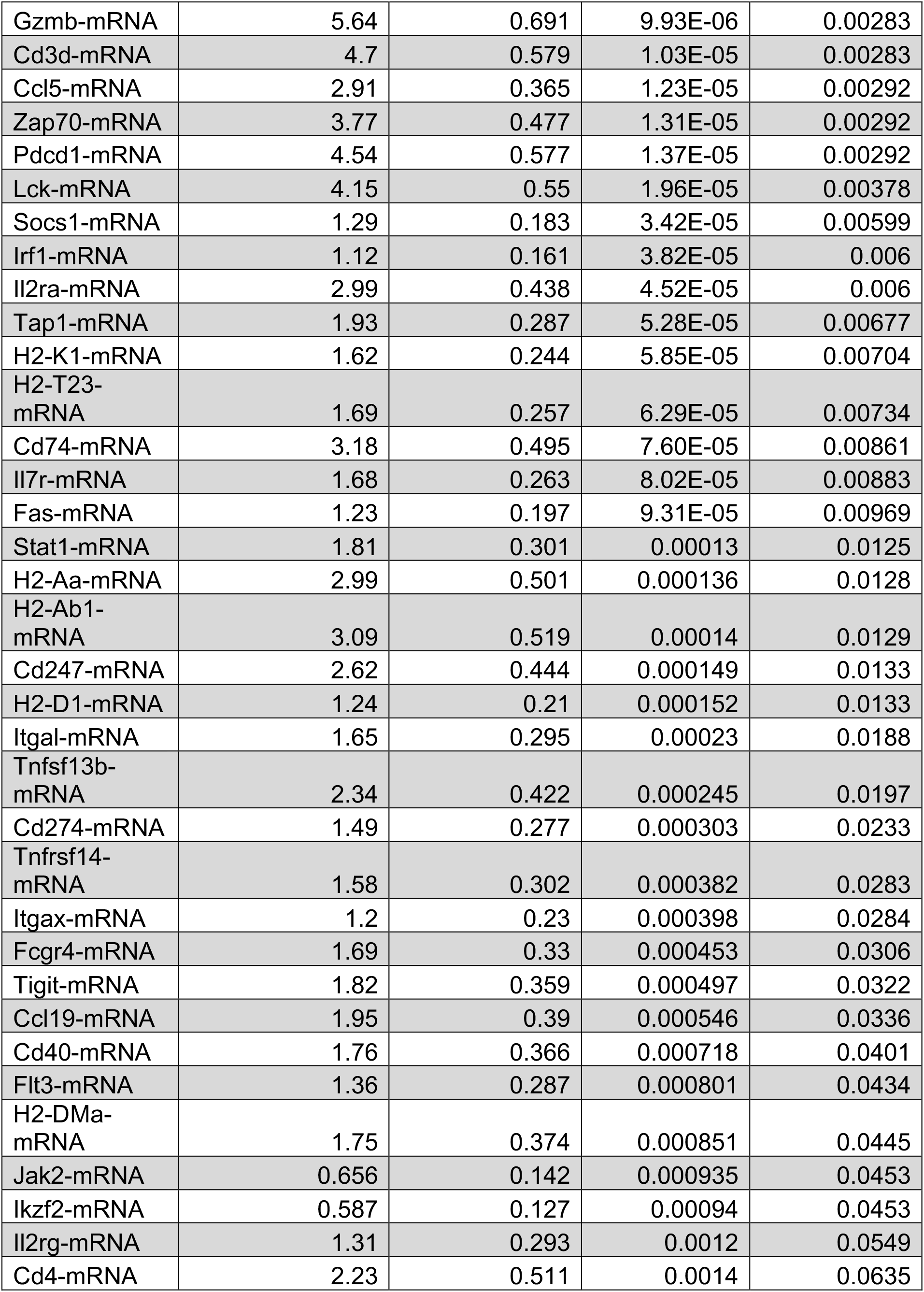
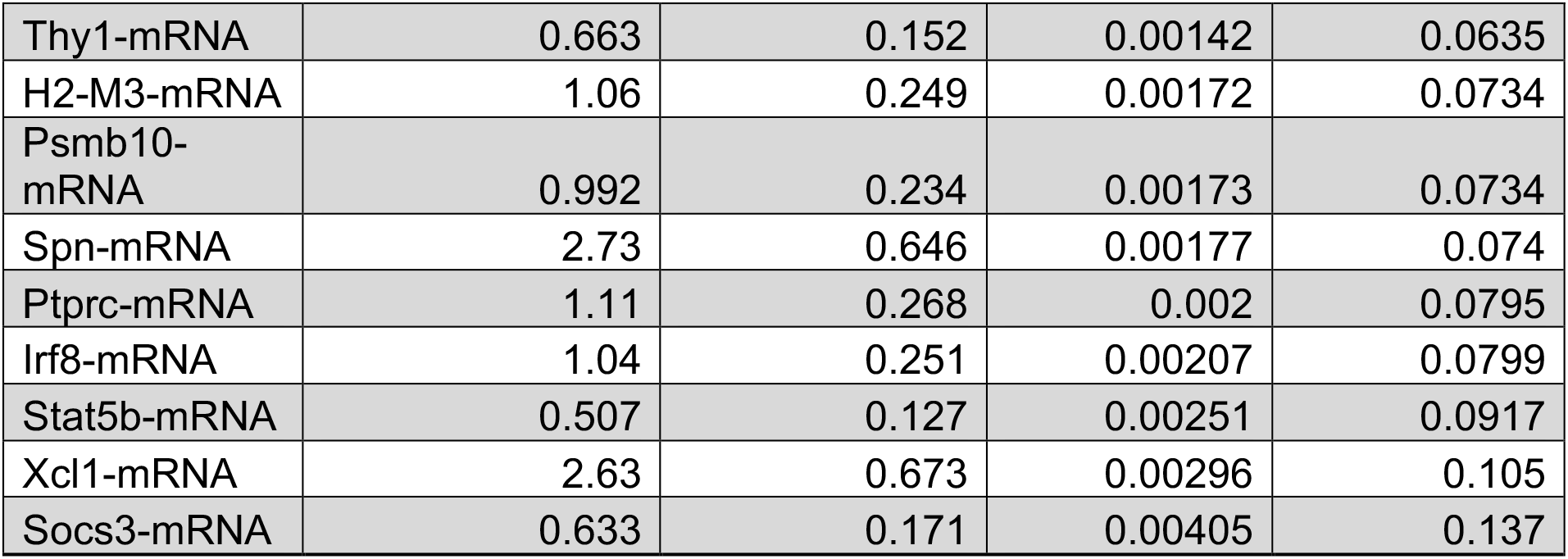
Differentially expressed T cell pathway genes.

**Supplementary Table 3:**
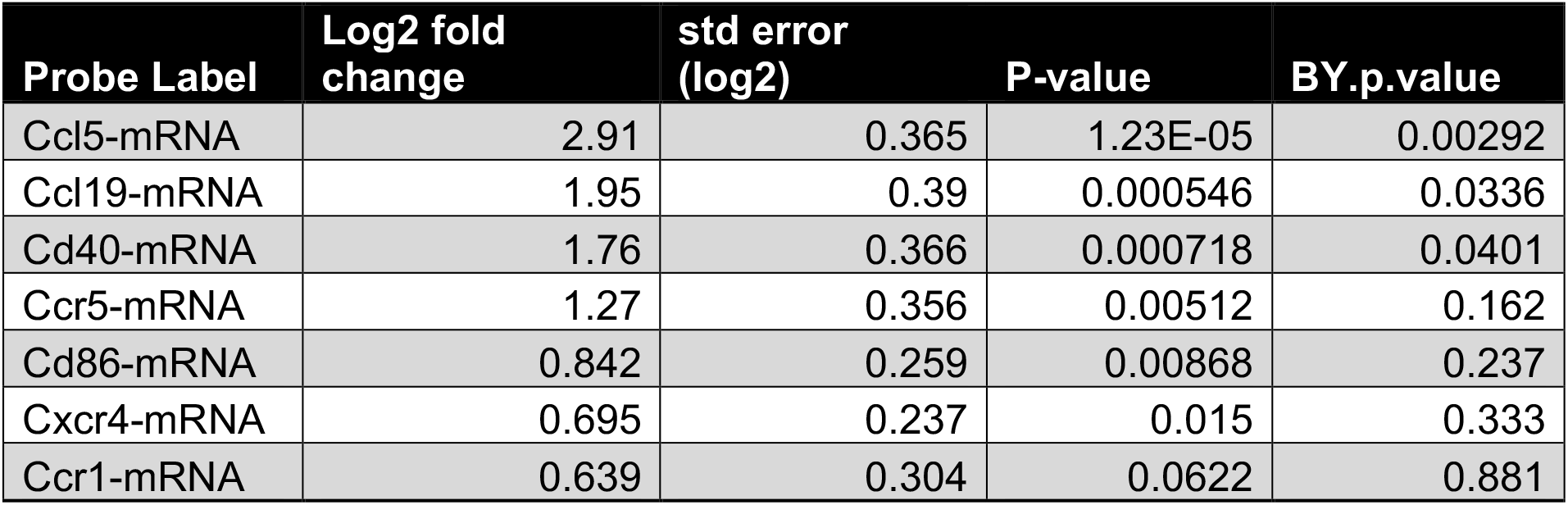
Differentially expressed DC functions pathway genes.

**Supplementary Table 4:**
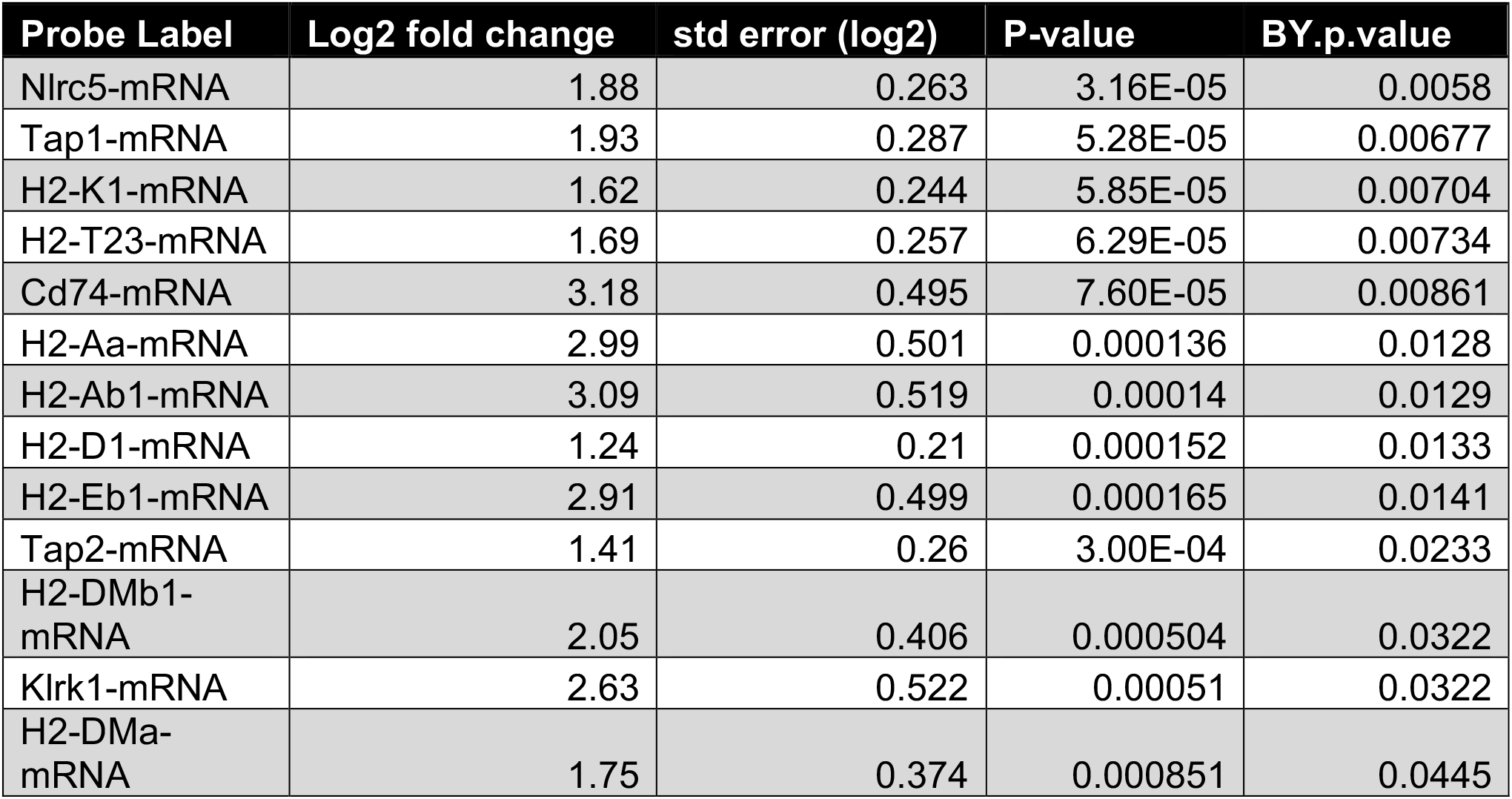
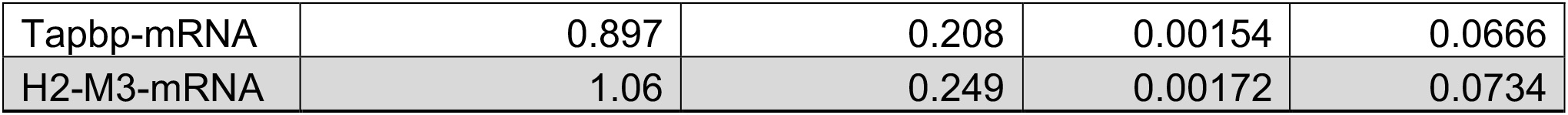
Differentially expressed MHC pathway genes.

